# Introducing double bouquet cells into a modular cortical associative memory model

**DOI:** 10.1101/462010

**Authors:** Nikolaos Chrysanthidis, Florian Fiebig, Anders Lansner

## Abstract

We present an electrophysiological model of double bouquet cells and integrate them into an established cortical columnar microcircuit model that has previously been used as a spiking attractor model for memory. Learning in that model relies on a Bayesian-Hebbian learning rule to condition recurrent connectivity between pyramidal cells. We here demonstrate that the inclusion of a biophysically plausible double bouquet cell model can solve earlier concerns about learning rules that simultaneously learn excitation and inhibition and might thus violate Dale’s Principle.

We show that learning ability and resulting effective connectivity between functional columns of previous network models is preserved when pyramidal synapses onto double-bouquet cells are plastic under the same Hebbian-Bayesian learning rule. The proposed architecture draws on experimental evidence on double bouquet cells and effectively solves the problem of duplexed learning of inhibition and excitation by replacing recurrent inhibition between pyramidal cells in functional columns of different stimulus selectivity with a plastic disynaptic pathway. We thus show that the resulting change to the microcircuit architecture improves the model’s biological plausibility without otherwise impacting the models spiking activity, basic operation, and learning abilities.

## 1 Introduction

We examine and build on a cortical microcircuit model, previously used in a working memory model by Fiebig and Lansner (2017) that implemented a BCPNN (Bayesian Confidence Propagation Neural Network) learning rule. We then expand on this functional columnar architecture by integrating GABAergic double bouquet cells (DBCs), which may play a key modulatory role in the cortical microcircuit (Krimer et al., 2005; Kelsom and Lu, 2013). Generally speaking, the BCPNN learning rule processes spike trains of pre- and postsynaptic neurons, and computes synaptic traces of activation and coactivation, which are then used to calculate the updated weights (see Sect. 2.3). In other words, the BCPNN is based on spike train correlations (Tully et al., 2014).

The foremost points of this work are first, the electrophysiological modeling of the inhibitory DBCs, and secondly their integration with the previous cortical memory model and its learning rule, which yields a novel model with improved biological plausibility and maintained functionality. The previous implementation suffers from the problem that learned weights among excitatory pyramidal cells in competing functional columns become negative, thus violating Dale’s Principle which states that neurons release the same neurotransmitters at all of their synapses (Strata and Harvey, 1999). The biological plausibility of our model is improved by a disynaptic interpretation of learned inhibition, whereby DBCs mediate this inhibition among pyramidal cells (Silberberg and Markram, 2007).

## 2 Materials and Methods

### 2.1 Neuron Models

We use an AdEx IAF (Adaptive Exponential integrate-and-fire) neuron model with spike-frequency adaptation. The neuron model has been modified for compatibility with a BCPNN synapse model (Tully et al., 2014) and reparameterized for simulation of several different neuron types. The model describes the temporal development of the membrane potenial *V*_m_ and the adaptation current *I*_w_, given by the following equations:

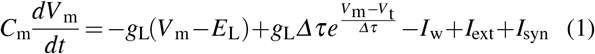

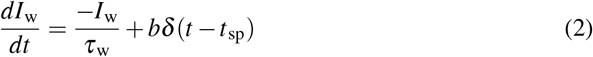

*Vm* represents cell membrane potential, *Iw* stands for the adaptation current, *Cm* is the membrane capacitance, g_L_ is the beak conductance, E_L_ is the leak reversal potential, V_t_ is the spiking threshold, *Δ_τ_* is the spike slope factor, b is the spike-triggered adaptation and *τ_w_* is the adaptation recovery time constant. Besides the stimulation current *I_ext_*, neurons receive synaptic currents *I_synj_* from AMPA and GABA synapses summed at the membrane:

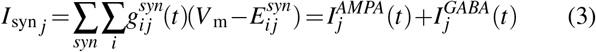

Adaptation enriches neural dynamics particularly particularly in pyramidal cells (Brette and Gerstner, 2005), and we take advantage of this in modeling DBCs as well.

### 2.2 Synapse model

The model features conductance based AMPA (reversal potential *E*^AMPA^) and GABA (reversal potential E^GABA^) synapses.

Plastic AMPA synapses under the spike-based BCPNN learning rule, are also subject to synaptic depression following the Tsodyks-Markram formalism (Tsodyks and Markram, 1997):

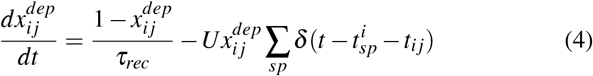

### 2.3 BCPNN learning rule

AMPA weights develop according to the BCPNN learning rule. BCPNN calculates three synaptic memory traces, *P_i_*, *P_j_* and *P_i j_*, implemented as exponentially weighted moving averages of pre-, post- and co-activation. As old memories deteriorate they are gradually replaced by newly learned patterns, so exponentially moving averages prioritize recent patterns. Specifically, BCPNN implements a three-stage procedure of exponential filters which defines Z, E and P traces. The method then estimates *P_i_* (normalized presynaptic firing rate), *P_j_* (normalized postsynaptic firing rate) and also *P_i j_* (coactivation) from these traces. In the final stage, *P_i_*, *P_j_* and *P_i j_* update the Bayesian weights *w_ij_* and biases β_*j*_. It is worth adding that E traces that enable delayed reward learning, are not used here because such conditions are not applicable. Some of the key equations are highlighted in this chapter; yet for further information and deeper understanding of the BCPNN learning rule, see Tully et al. (2014).

To begin with, BCPNN receives pre- and postsynaptic spike trains (*S_i_*,*S _j_*) so as to calculate the traces *Z_i_* and *Z_j_*:

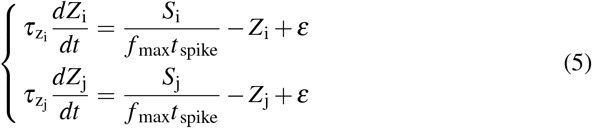

*f*_max_ denotes the maximal neuronal spike rate, ε is the lowest attainable probability estimate, *t_spike_* denotes the spike duration while τ_z_i = τ_z_j are the pre- and postsynaptic time constants respectively (here 5 *ms*).

P traces then are estimated from the Z traces as follows:

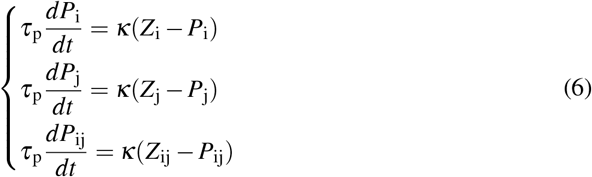

The parameter κ adjusts learning speed, and by setting κ = 0 there are no weight changes. To give prominence to the stability of memory networks with BCPNN learning rule, we set κ = 1 during the whole simulation.

Finally, *P_i_*, *P_j_* and *P_ij_* are used to calculate intrinsic excitability *β*_j_ and synaptic weights *w_i j_*:

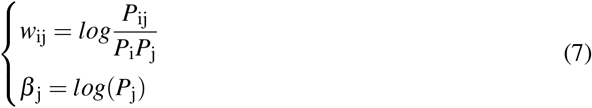

### 2.4 Simulation tools

We use NEST (Neural Simulation Tool) version 2.4.2, and a custom-built BCPNN learning rule module (Tully et al., 2014). NEST simulates the dynamics of spiking neural models and features a convenient Python interface (PyNEST), providing access to NEST’s simulation kernel (Gewaltig and Diesmann, 2007).

## 3 Results

### 3.1 Double bouquet cells

Modeling DBCs is a key contribution of this work since the suggested cortical microcircuit model learns disynaptic inhibition through them and thereby modulates the neural activity of neurons in competing MCs. DBCs are GABAergic interneurons which may play an important role in shaping neural activity and circuitry (Krimer et al., 2005; Kelsom and Lu, 2013) and are mainly located in Layer II/III featuring a bitufted dendritic conformation (Markram et al., 2004). They contact the dendrites of targeted cells (Markram et al., 2004) innervating spines (69.2% *±* 4.2%) and shafts (30.8% ± 4.2%) (Tamas et al., 1997). DBCs are characterized by vertically oriented descending axons (María and DeFelipe, 1995), which are generally termed “bundles” or “horse-tails” (Yáñez., 2005; DeFelipe et al., 1989).

The majority of double bouquet cells appear to be situated in upper layers (DeFelipe et al., 1989) and one of their unique feature is a horse-tail that fit well within the minicolumn (vertical cyclinder of tissue with a diameter of roughly 25−50 μ*m*). Due to this morphology they create strong connections with pyramidal cells within their local column (DeFelipe et al., 2006). Neuroanatomical data suggests that each minicolumn contains one DBC (DeFelipe et al., 2006).

DBCs present a small degree of adaptation (Tamas et al., 1997) and unlike Basket Cells, they are characterized by sustained spiking activity (Krimer et al., 2005), which under strong stimulation is at 43 ± 13 *Hz* (Zaitsev et al., 2008). DBCs are intermediate spiking (IS) cells (Krimer et al., 2005) and subclassified as RSNP cells (Regular-spiking nonpyramical neurons) according to studies of their physiological properties (Kawaguchi and Kubota, 1996, 1997).

*In vitro* experiments report DBC resting potential at *−*76 *±* 6 *mV*, spiking threshold *V*_th_ at *−*44 *±* 8 *mV* and input resistance at 626 *±* 312 *MΩ*. They are considered inhibitory neurons with high input resistance (Zaitsev et al., 2008) and low capacitance (see Table 1). In addition, they exhibit membrane time constants (17.1 7.7 *ms*), slope factor (0.64 2.22) and action potential amplitude (66 12.4 *mV*) (Krimer et al., 2005).

**Table 1.** DBC model parameters

| Parameters | Symbol | Value |
| --- | --- | --- |
| Adaptation current | $b$ | 3 pA |
| Adaptation time constant | $\tau_w$ | 200 ms |
| Membrane capacitance | $C_m$ | 15 pF |
| Leak reversal potential | $E_L$ | -76 mV |
| Leak conductance | $g_L$ | 1.52 pS |
| Upstroke slope factor | $\Delta_t$ | 3 mV |
| Spike threshold | $V_t$ | -44 mV |
| Spike reset potential | $V_r$ | -62 mV |
| Refractory period | $\tau_{ref}$ | 2 ms |
| Input Resistance | $RI$ | 660 MΩ |

We align simulation model for DBCs with biological findings, yet tune some factors such as adaptation (*b*), leak conductance (*g*_L_), slope factor (Δ _t_) and refractory period (τ_ref_) to achieve satisfactory electrophysiological fidelity, reproducing spike patterns under sweeps of increasing suprathreshold current steps and other typically reported activity. Fig. 1C displays the membrane voltage of a stimulated DBC. The resulting model parameters are broadly consistent with experimentally reported values (see Table 1).

**Fig. 1.**
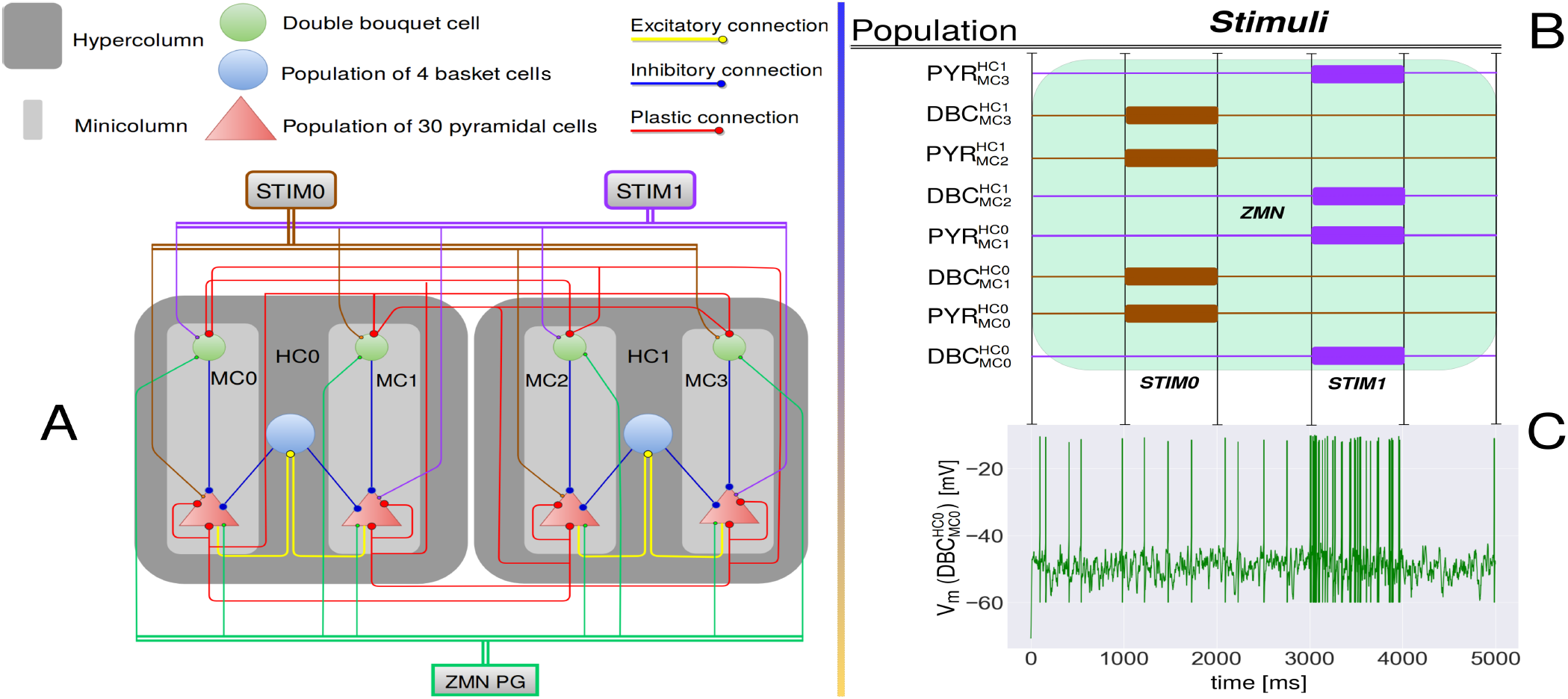
**A)** The proposed functional columnar architecture including DBCs. The model contains 2 Hypercolumns (HC0,HC1) nesting 2 mini-columns each (MC0, MC1, MC2, MC3). Each MC contains one DBC, which delivers inhibition to all pyramidal cells in their MC. MCs are selective for different stimuli such that MC0 and MC2 function coactively and compete with another pair of coactive minicolumns MC1 and MC3. When MC0 and MC2 are active, MC1 and MC3 are silent and vice versa. Pyramidal neurons of MC0 and MC2 disynaptically inhibit MC1 and MC3 through basket cells (within HCs) and DBCs (within and between HCs). Plastic connections are drawn with 20% connection probability, while the connections between basket and pyramidal cells within the local hypercolumns are drawn with 70% connection probability. Conductance delays within and between HCs are 1.5 and 4.5 *ms* respectively. **B)** Training stimuli drive pyramidal and DBCs. 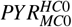 denotes the pyramidal cells in MC0. A poisson generator (STIM0 - brown) stimulates 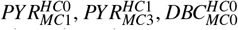 and 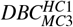 between 1000 *ms* and 2000 *ms*. Another poisson generator (STIM1 - purple) stimulates 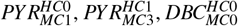 and 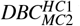 between 3000 *ms* and 4000 *ms*. A zero mean noise poison generator (ZMN - green shaded area) is active throughout the simulation. **C)** Membrane voltage of a stimulated DBC. STIM1 specifically drives this cell between 3000 *ms* and 4000 *ms* (cf. Fig. 1B). The DBC presents sustained low-rate firing throughout simulation and reaches typically reported firing rates during stimulation (see Table 1).

### 3.2 Adding DBCs to the columnar architecture

The proposed architecture principally follows several previous spiking neural network implementations (Lansner, 2009; Fiebig and Lansner, 2017) and is best understood as a subsampled cortical layer II/III model with nested hyper-columns (HCs) and minicolumns (MCs); yet in this new model, connections among pyramidal cells in competing MCs are mediated by DBCs as an additional local microcircuit component (see Fig. 1A). Their functional role is to deliver the same amount of inhibition to the respective MCs as the previous model but now entirely disynaptically, without principally changing established network dynamics.

The thereby extended cortical microcircuit model now contains three classes of neurons; pyramidal cells, basket cells and DBCs (see Fig. 1A). We use parameters for pyramidal and basket cells from a previous model implementation (Fiebig and Lansner, 2017) and derive parameters for DBCs through electrophysiological modeling and tuning based on reported *in vitro* characterizations (see Sect. 3.1). We simulate two reduced HCs compared to previous larger network models.

### 3.3 BCPNN plasticity

Stimulation of the columnar network changes the efficacy of the plastic BCPNN synapses. To show the learned connectivity we read out connection weights after an one second long initialization with zero mean noise (Initial Weight Distribution, IWD) and finally, after learning of the two stimuli (Learned Weight Distribution, LWD), see Fig. 1B.

Fig. 2 shows histograms of plastic weights between the pyramidal cells in MC0 and their post-synaptic targets. Fig. 2A and 2B show that the integrated DBCs in the columnar architecture do not affect the behaviour of the functional weights which are not related to them.

**Fig. 2.**
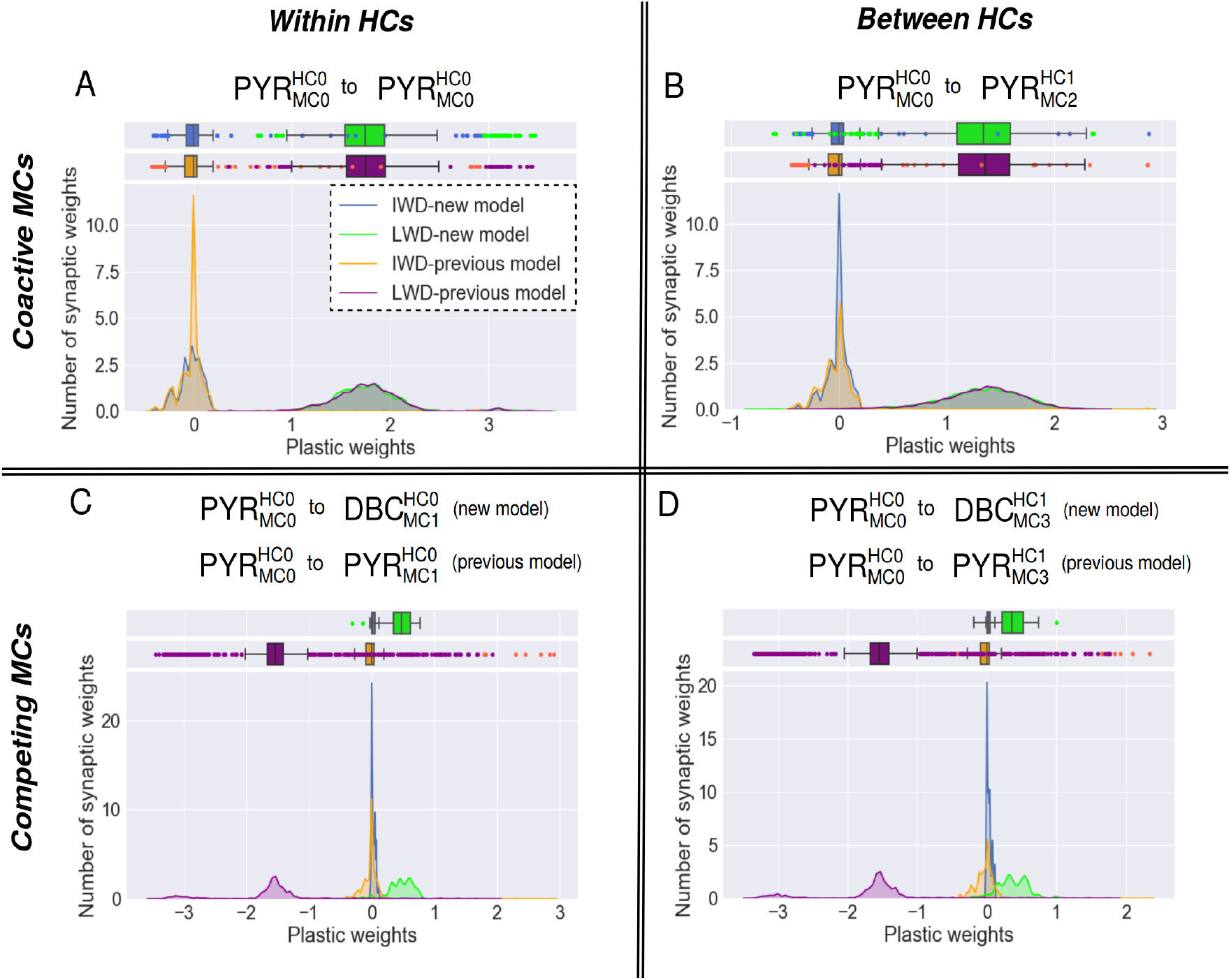
Weights distribution of plastic connections from pyramidal cells in MC0 before (Initial Weight Distribution, IWD) and after (Learned Weight Distribution, LWD) training in the previous and updated microcurcuit (multi-trial averages). IWD (new model) in blue, LWD (new model) in green, IWD (previous model) in orange and LWD (previous model) in purple. **A)** Recurrent AMPA weight distribution among the pyramidal cells of MC0. Initial and converged distributions from both columnar networks are similar with means of 0 and 1.7 respectively, (see plot-box). **B)** Distribution of AMPA weights between the pyramidal cells of the coactive MC0 and MC2 (assosiate connection between HCs). The distribution of the initial weights has the same behavior in both architectures, with a mean close to 0. The distributions of the learned weights overlap, with a mean of 1.3. **C)** The initial distribution of plastic weights in both functional columnar architectures is close to 0. The negative learned weights of the previous model (violating Dale’s principle) are functionally substituted in the new model by positive weights from the pyramidal cells of MC0 to the DBC of the competing MC1. The converged positive weight distribution has a mean of 0.5. **D)** Plastic connections between pyramidal cells of MC0 and DBC of the competing MC3 are compared with plastic connections between pyramidal cells in MC0 and MC3 in the previous model. The behaviour remains the same as in C) because the negative weights turn positive after DBCs integration.

Two important changes can be identified in Fig. 2C and 2D, wherein disynaptic inhibition is introduced by DBCs. Even though the strength of the learned connections onto DBCs is weak, DBCs also feature low capacitance and dense connections with local pyramidal cells, and thus deliver comparable inhibition (see Sect. 3.4).

This result shows the effectiveness of DBCs involvement in the microcircuit network as they learn to mediate disynaptic inhibition between pyramidal cells in competing MCs. This outcome looks promising, but we yet have to verify its functional efficacy with regards to the total inhibition delivered (see Sect. 3.4).

### 3.4 Functionality verification

Fig. 3A displays the spiking activity of neurons in a simulated HC (HC0). Although DBCs keep a low level sustained spiking activity throughout the simulation, they can reach higher firing rate during training (Zaitsev et al., 2008).

The cortical model learns as expected and the competing MCs inhibit each other by disynaptic inhibition mediaded by DBCs and basket cells. But is this inhibition equivalent to the mono-synaptic inhibition learned by the previous model?

We tested learning in both the new and previous model using the same stimulation pattern and recorded the total inhibitory input current received by pyramidal cells in MC0 (see Fig. 3B). The proposed cortical model effectively delivers the same amount of disynaptic inhibition via basket cells and DBCs. The new model has thus improved biological credibility while maintaining the same functionality.

**Fig. 3.**
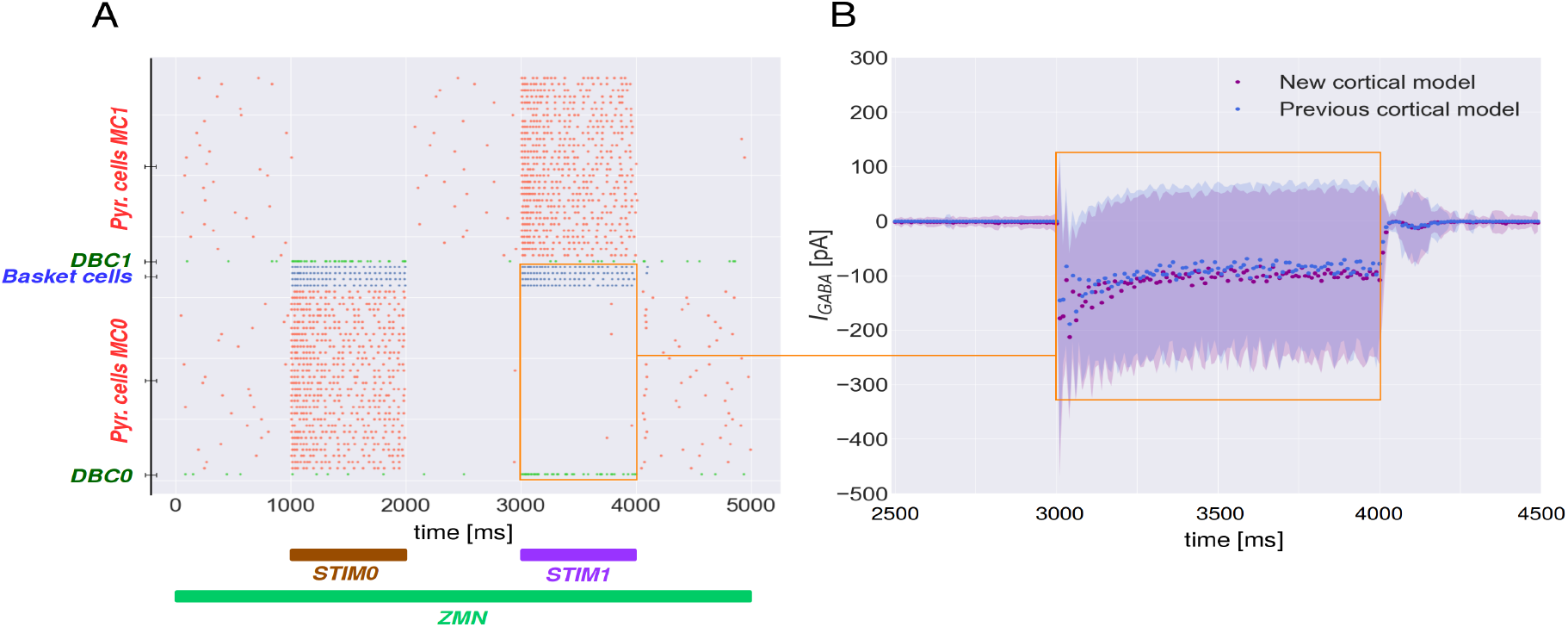
**A)** Spike raster of neurons in HC0. Colors correspond to Fig. 1A, 1B. When STIM0 or STIM1 are ON (cf. Fig. 1B), pyramidal cells in MC0 excite basket cells within their local MC and DBC in competing MC which in turn inhibit pyramidal cells in MC1 and vice versa. **B)** Population averaged total inhibitory input current received by pyramidal cells in MC0 in both architectures (100 trials, 10 *ms* bins). New model in purple, previous network in blue. DBC and basket cells simultaneously deliver inhibition to the neurons of MC0. The total inhibition current (*I*_GABA_) starts from zero level (2500 *ms*-3000 *ms*), then decreases (3000 *ms*-4000 *ms*) reaching a climax of 220 *pA* and finally stabilizes at zero (4000 *ms*-4500 *ms*) following the same pattern in the new and previous cortical model.

## 4 Discussion

This work aims at giving prominence to the double bouquet cells and their use as an integral part of a cortical microcircuit model. The population of DBCs is limited compared to other GABAergic neurons; however, they may play a key role in shaping neural activity. By integrating them into an established model, the new model now obeys Dale’s principle with maintained function. Indirectly, this result also verifies the biological plausability of recent network models (Lansner, 2009; Tully et al., 2016; Fiebig and Lansner, 2017). The newly integrated DBCs effectively learn to mediate disynaptic inhibition between pyramidal cells thus eliminating negative learned weights between pyramidal cells which violate Dale’s principle.

In conclusion, the successful integration of an electrophysiological DBC model into an established cortical microcircuit design yields a novel functionally equivalent learning network with improved biological plausibility. This model suggests that DBCs have a quite well defined role in cortical memory networks.

## Acknowledgements

Funding from T.E.I of Athens, Erasmus Placement [NC], EuroSPIN Erasmus Mundus Doctoral Programme [FF], and Swedish E-Science Research Center (SeRC) [AL] is gratefully acknowledged.

